## Supplemental Figure and Table 1 for "Rediscovering the *Rete Ovarii*: a secreting auxiliary structure to the ovary"

### SUPPLEMENTAL INFORMATION

### Supplemental Figure 1

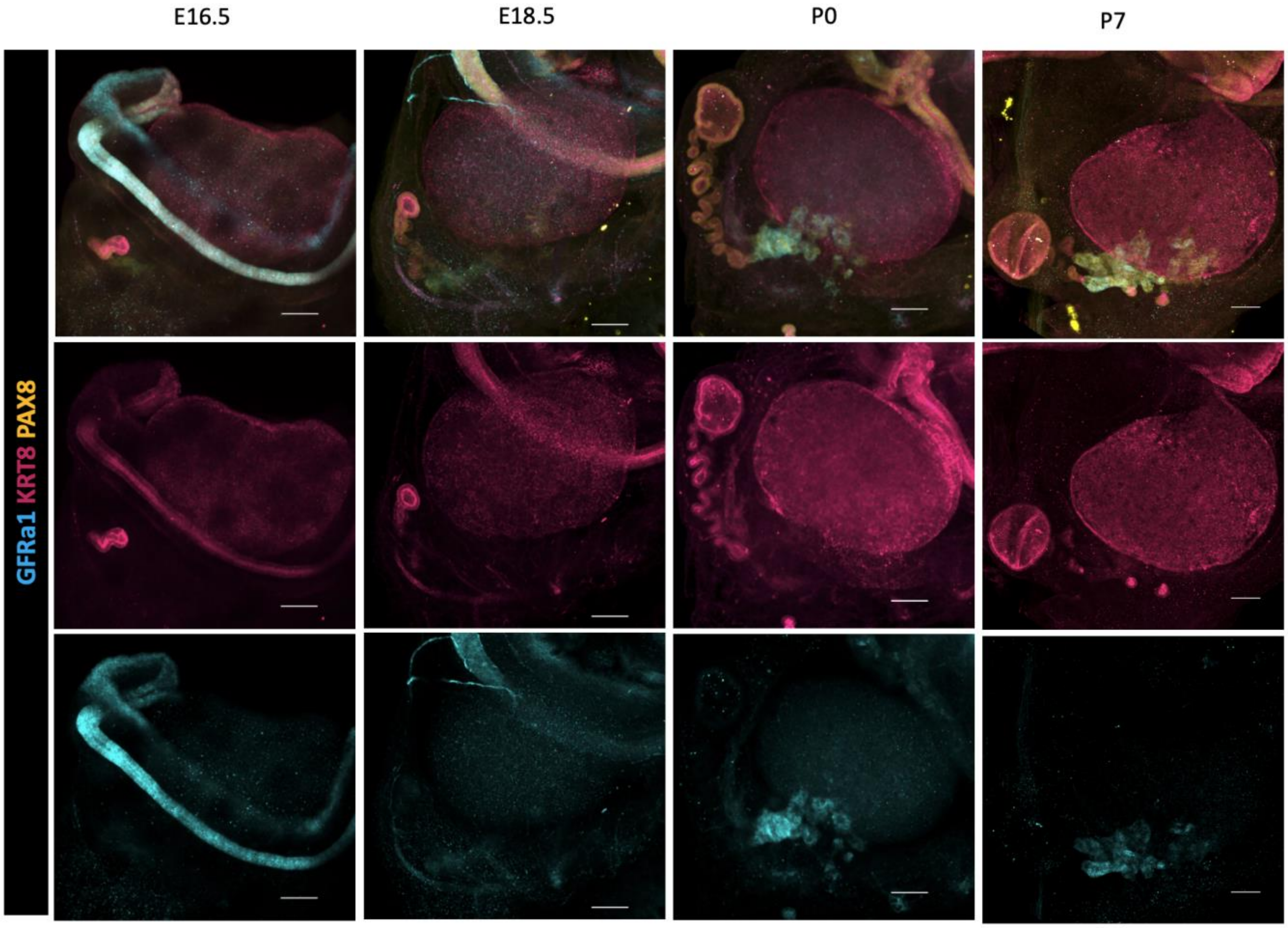


*Supplemental Figure 1. Ventral view of ovary and EOR during development.*

*Maximum intensity projection from confocal Z-stacks of whole ovary/mesonephros complexes at E16.5 (a), E18.5 (b), P0 (c) and P7 (d) immunostained for PAX8 (yellow), GFRa1 (cyan) and KRT8 (magenta). Top row are composite images, while the third row show separate panels for KRT8 and and the bottom row show panels for GFRa1. *Note that GFRa1 and KRT8 do not co-localize and are specific to the CR and EOR, respectively. All figures are dorsal views of the ovary. Yellow asterisk indicates opening of the infundibulum for reference. Scale bar – 100um*

### Supplemental Table 1

| Gene Name | Sequence 5' to 3' |
|---|---|
| mTmG (Mutant Reverse) | AAT CCA TCT TGT TCA ATG GCC GAT C |
| mTmG (Wildtype Forward) | GGC TTA AAG GCT AAC CTG ATG TG |
| mTmG (Mutant Forward) | CCG GAT TGA TGG TAG TGG TC |
| mTmG (Wildtype Reverse) | GGA GCG GGA GAA ATG GAT ATG |
| Tre-Cre (Forward) | GCG GTC TGG CAG TAA AAA CTA TC |
| Tre-Cre (Reverse) | GTG AAA CAG CAT TGC TGT CAC TT |
| Pax8-rtTa (Forward) | CCA TGT CTA GAC TGG ACA AGA |
| Pax8-rtTa (Reverse) | CTC CAG GCC ACA TAT GAT TAG |
| Tre-H2B-GFP (Forward) | GCT CGT TTA GTG AAC CGT CAG |
| Tre-H2B-GFP (Reverse) | TCT TCT GCG CCT TAG TCA CC |
| LGR5 (Mutant Forward) | CTG AAC TTG TGG CCG TTT AC |
| LGR5 (Wildtype Reverse) | GTC TGG TCA GAA TGC CCT TG |
| LGR5 (Common) | CTG CTC TCT GCT CCC AGT CT |

*Supplemental Table 1. Table recapitulating the DNA forward and reverse primers used for genotyping the transgenic mouse lines used in the present study*

### Supplemental movie legend

**Supplemental Movie 1. 3D model of an XX ovarian complex at E18.5**

This video depicts the 3D rendering of native lightsheet images of an E18.5 ovary/mesonephros complex. Demonstrating how the rete wraps around the ovary. DNA (grey) Ovarian Surface epithelium (green) PAX8 (red).
